# Perilipin membrane-integration determines lipid droplet heterogeneity in differentiating adipocytes

**DOI:** 10.1101/2023.10.30.564726

**Authors:** Mario Majchrzak, Ozren Stojanović, Dalila Ajjaji, Kalthoum Ben M’barek, Mohyeddine Omrane, Abdou Rachid Thiam, Robin W. Klemm

**Affiliations:** Department of Physiology Anatomy and Genetics, University of Oxford, Oxford OX1 3PT, UK; Department of Molecular Life Sciences, University of Zurich, 8057 Zurich, CH; Laboratoire de Physique de l’École Normale Supérieure, ENS, Université PSL, CNRS, Sorbonne Université, Université de Paris, F-75005 Paris, France

**Keywords:** LD-heterogeneity, LD-targeting, ERTOLD, CYTOLD, LD-affinity, PLIN1

## Abstract

The storage of fat within lipid droplets (LDs) of adipocytes is critical for whole-body health. Acute fatty acid (FA) uptake by differentiating adipocytes leads to the formation of at least two LD-classes marked by distinct perilipins (PLINs). How this LD-heterogeneity arises is an important yet unresolved cell biological problem. Here, we show that an unconventional integral membrane-segment (iMS) targets the adipocyte specific LD-surface factor PLIN1 to the endoplasmic reticulum (ER) and facilitates high affinity binding to the first LD-class. The other PLINs remain largely excluded from these LDs until FA-influx recruits them to a second LD-population. Preventing ER-targeting turns PLIN1 into a soluble, cytoplasmic LD-protein, reduces its LD-affinity and switches its LD-class specificity. Conversely, moving the iMS to PLIN2 leads to ER-insertion and formation of a separate LD-class. Our results shed light on how differences in organelle targeting and disparities in lipid-affinity of LD-surface factors contribute to formation of LD-heterogeneity.

**Highlights:**

- PLIN1 is an integral membrane protein behaving as a class I LD protein.
- An unconventional integral membrane segment (iMS) mediates ER insertion.
- The iMS is required for LD-heterogeneity in differentiating adipocytes.
- High affinity LD-targeting from the ER is likely a gated process.

**eTOC blurb:** Majchrzak et al. use biochemistry, imaging and *in vitro* experiments to identify molecular features in PLIN1 that determine ER insertion and high lipid droplet affinity which are both important in the generation of LD-heterogeneity within differentiating adipocytes.

## Introduction

Adipocytes deposit an excess of dietary nutrients as triacylglycerols (TAGs) in large lipid droplets (LDs) and release fatty acids to feed other tissues during fasting^1–3^. Deficiencies in LD biogenesis and turnover are associated with several metabolic diseases such as lipodystrophies, fatty liver disease, type-2 diabetes, and obesity^4–6^.

LDs form at the membranes of the endoplasmic reticulum (ER) when neutral lipids, such as TAGs, accumulate between the two leaflets of the lipid bilayer^7–9^. When TAG-amounts increase they phase-separate and form lenses, which eventually bud out into the cytoplasm as LDs. The LD-surface is surrounded by a phospholipid monolayer and specific surface proteins^10–15^.

In principle all LDs within a cell should have the same lipid and protein composition. Accumulating evidence suggests, however, that LDs are highly heterogenous, and can specialize into different sub-populations which are morphologically, chemically, and functionally distinct^14,16–22^.

One way by which cells can generate different LD-classes is through the synthesis of the hydrophobic core. For example, fly fat bodies contain a population of large LDs, which is mostly filled with TAGs derived from *de novo* lipogenesis^23,24^. A class of smaller LDs in the periphery of the cell seems to store TAGs made preferentially from dietary fatty acids (FAs)^23,24^. Consistent with these findings, FA feeding of mammalian adipocytes leads to the formation of distinct LD-classes^10,25,26^. A central LD population is formed mainly by *de novo* lipogenesis^27^, whereas acute FA influx triggers the formation of a second, peripheral LD class^10,25,26^. The mechanisms that lead to the formation of different LD populations remain, however, poorly understood, and it is largely unclear how LD-surface factors can be targeted to separate LD-populations in the cell.

In general, protein targeting to LDs occurs by two major pathways^14,28,29^. Class I proteins are synthesized as monotopic integral membrane proteins at the ER, and subsequently move to the LD surface using the ERTOLD pathway (**Fig. 1A**, class I). The dual localization to the ER and the LDs is facilitated by special membrane anchors that often fold into hairpins^13,29^. Class II (or CYTOLD) proteins, on the other hand, bind LDs directly from the cytoplasm (**Fig. 1A**, class II). They do not possess integral membrane segments and are instead targeted to the LD by amphipathic domains detecting packing defects in the LD monolayer where the hydrophobic LD core is exposed^14,30–35^.

**Figure 1.**
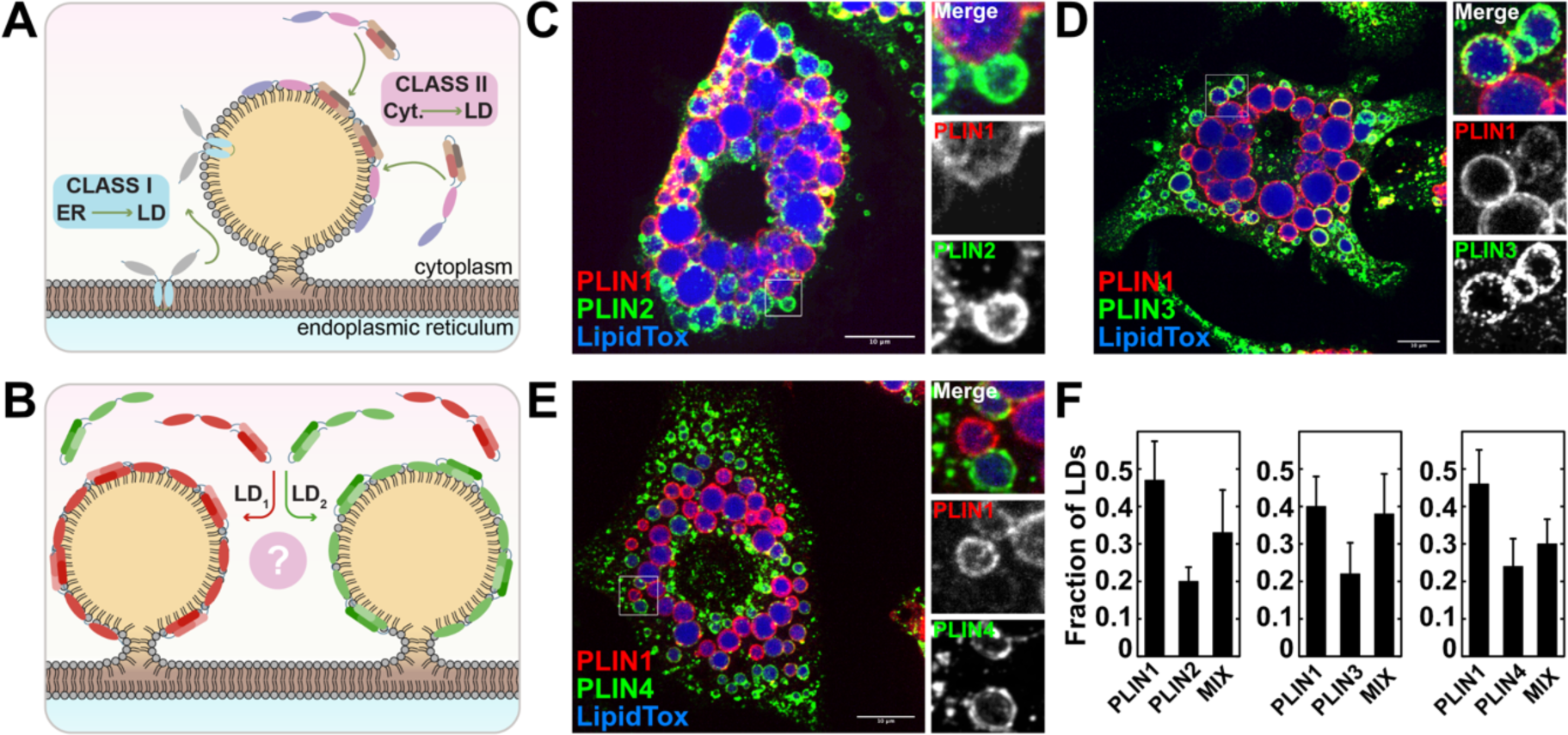
Fatty acid feeding leads to the formation of at least two distinct LD-classes in adipocytes. **(A)** Class-I LD proteins move from the ER to the LD surface. Class-II LD proteins bind to LDs directly from the cytoplasm. **(B)** Segregation of class-II proteins to different LD-classes. **(C)** Day 6 3T3-L1 adipocytes fed with 750µM Na^2+^oleate for 3h. Endogenously expressed PLIN1 and PLIN2 were visualized by immunofluorescence. Scale bar, 10 µm. **(D)** As in (C) but PLIN1 and PLIN3 were imaged. **(E)** As in (C) but PLIN1 and PLIN4 were imaged. **(F)** Quantification of the results in (C)-(E). n=9; n=9; n=8, for cells co-stained with PLIN1 and 2 or PLIN1 and 3 or PLIN1 and 4, respectively. LD counts per replicate >100; values are expressed as mean LD in each class ± SEM.

The perilipins (PLINs) are currently viewed as prototypical class II proteins, comprising a family of evolutionarily conserved LD-surface factors carrying out important structural and regulatory functions^26,36–39^. The family defining amphipathic PAT domain and the following 11-mers mediate LD-targeting (**Fig. S1A**)^32,37,40,41^. The PLINs are all thought to target the LD from the cytoplasm and should therefore mix on the same LD class. The LD sub-populations that form in differentiating adipocytes upon FA feeding are, however, marked by specific PLINs (**Fig. 1B**) ^10,25,26^. An affinity-based LD binding “hierarchy” among the different PLIN-family members is likely key to the formation of the two LD classes^34,40,42–45^. However, the biochemical characterization of the responsible targeting domains remains incomplete^38^.

## Results

### Acute fatty acid feeding induces the formation of distinct LDs in adipocytes

To investigate perilipin (PLIN) targeting to LDs we differentiated 3T3-L1 cells into white adipocytes and imaged endogenously expressed PLIN1, 2, 3, and 4 after FA feeding (PLIN5 is not expressed in 3T3-L1 adipocytes ^11,27^). Confirming previous observations^26,36,39,40,42^, we observed distinct LD populations (**Fig. 1B-F**). A central LD class was almost exclusively labelled with PLIN1, and a second, peripheral population was marked by PLIN2, 3, and 4 (Fig. 1C, D, and E). The second population was either exclusively marked by PLIN2, 3, and 4, or contained sometimes low amounts of PLIN1 which we called mixed LDs (**Fig. 1C-E** (zoom-ins), **F** (MIX)).

Consistent with our results in adipocytes and the established LD binding hierarchy^26,43^, we next showed that ectopically expressed PLIN1 excluded PLIN2, 3, or 4 from LDs when co-transfected in COS7 cells (**Fig. S1A-D**). This behaviour was dependent on features C-terminally of the 11-mer domain (**Fig. S1A**). A construct only consisting of the N-terminal PAT and 11-mer regions (PLIN1 1-192) mixed well with PLIN2 on the LD surface (**Fig. S1E**).

Together, these data raised the question of how class-II LD proteins can separate onto different LD-populations.

### PLIN1 is an integral membrane protein dually localizing to the ER and LDs

To determine the molecular basis of PLIN LD targeting, we expressed PLIN1 and 2 in COS7 cells. As expected, PLIN1 localized to the endogenous LDs but was, surprisingly, never seen in the cytoplasm. Instead, it always bound to the ER (**Fig. 2A, Fig. S2B**). Consistent with previous work^45^, these data indicated that PLIN1 is not a prototypical class-II LD protein. On the other hand, PLIN2, showed normal class-II behaviour accumulating in the cytoplasm upon overexpression and covering all endogenous LDs (**Fig. 2B, S2A, S2D**). After addition of FA (Na^+^ oleate), both PLIN1 and PLIN2 targeted efficiently to LDs (**Fig. 2C-E**).

**Figure 2.**
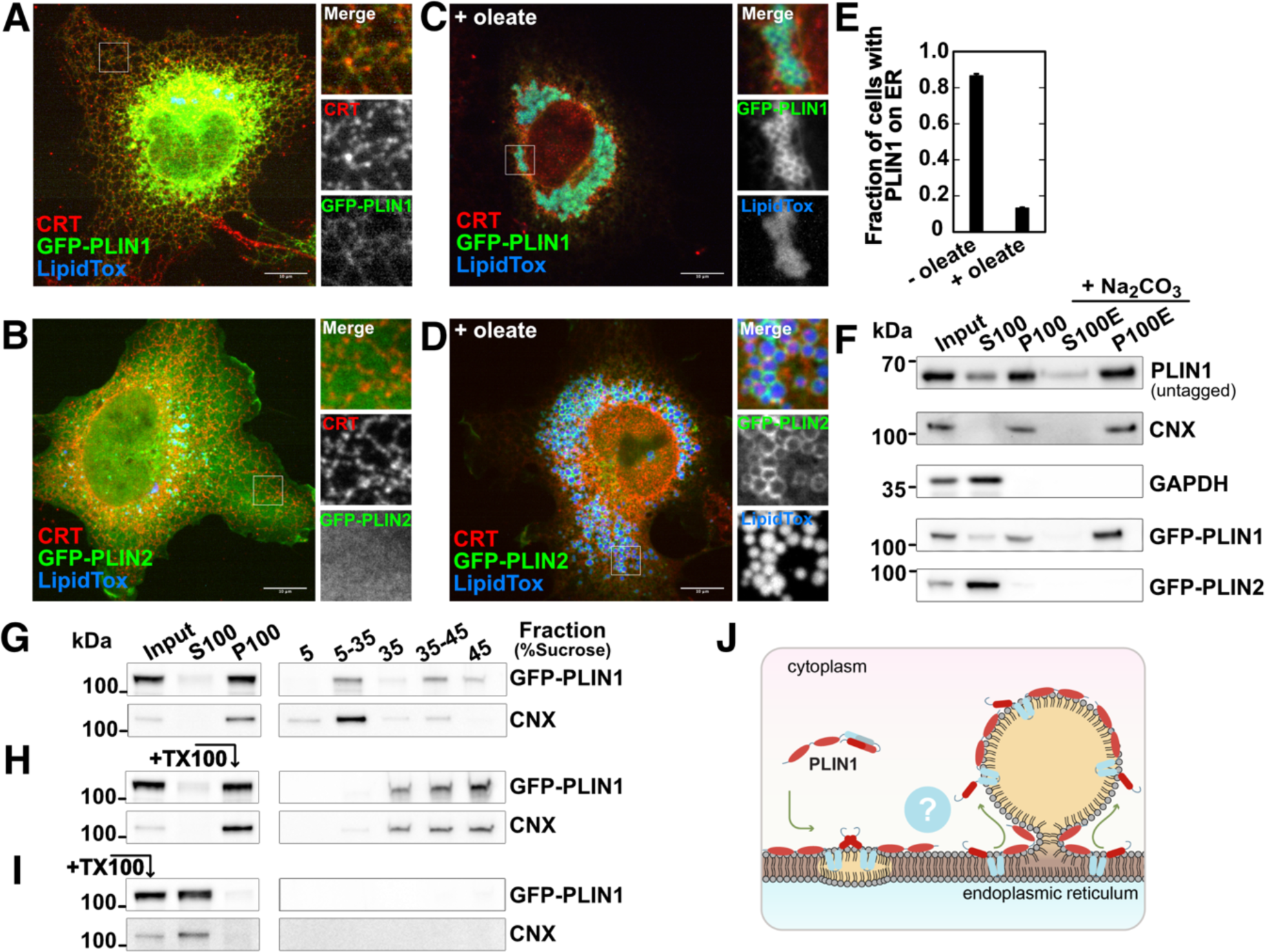
PLIN1 is an integral membrane protein which localizes to the ER and LDs. **(A)** GFP-murinePLIN1 was expressed in COS7 cells. The ER is visualized with an antibody against calreticulin (CRT). LDs are stained with LipidToxRed. Scale bar, 10 µm. **(B)** As in (A) but the cells expressed GFP-PLIN2. **(C)** As in (A) but with 250 µM Na^+^oleate for 6h. **(D)** As in (C) but the cells expressed GFP-PLIN2. **(E)** Quantification of results in (A) and (C) with GFP-PLIN1 on the ER. n = 3 independent experiments with >1500 cell per replicate; fraction of PLIN1 on the ER ±SD. **(F)** COS7 expressing untagged PLIN1, GFP-PLIN1 or GFP-PLIN2, were fractionated followed by alkaline extraction. **Input,** cell lysate; **S100, P100,** supernatant and pellet of a 100k x g centrifugation, **S100E, P100E,** E denotes alkaline (Na_2_CO_3_) extraction. **GAPDH,** glycerolaldehyde-3-phosphate dehydrogenase; **CNX,** calnexin. **(G)** As in (F), but P100 was floated on a sucrose-step gradient. **(H)** As in (G), but P100 was incubated with Triton X-100 before flotation. **(I)** As in (H), but the cell lysate was treated with Triton X-100 before centrifugation. **(J)** PLIN1 is an integral membrane protein with a membrane anchor (blue) allowing ER and LD-targeting.

Since PLIN1 does not contain any predicted membrane segments (**Fig. S2D, S2E**), and exhibits similar overall hydrophobicity as PLIN2 (**Fig. S2F, S2G**), we initially expected it to bind the ER as a peripheral membrane protein perhaps through a receptor. In support of this view, PLIN1 sedimented together with the endogenously expressed integral ER protein calnexin (CNX) to the membrane pellet (**Fig. 2F**, PLIN1 untagged; GFP-PLIN1, P100).

However, when we subjected the membrane pellet (P100) to alkaline extraction with Na_2_CO_3_-buffer, PLIN1 remained in the alkaline resistant membrane fraction (**Fig. 2F**, P100E) behaving like an integral membrane protein. PLIN2 co-fractionated with the soluble cytoplasmic marker glyceraldehyde-3-phosphate dehydrogenase (GAPDH, **Fig. 2F**, S100).

To exclude possible artefacts, we carried out flotation experiments with a ribosome stripped membrane fraction from cells expressing PLIN1. **Fig. 2G** shows that PLIN1 moved efficiently out of the bottom fraction and floated on a sucrose step gradient together with ER membranes marked by CNX (5-35% sucrose interface). Addition of detergent to P100 prevented flotation (**Fig. 2H**), and PLIN1 and CNX both remained soluble when we mixed the cell lysate with detergent before high-speed centrifugation (**Fig. 2I**).

Taken together, our data indicated that PLIN1 does not belong to the cytoplasmic class-II proteins. As an integral membrane proteins LD-targeting might move from the ER to the LD surface. Indeed, we found that PLIN1 ER-pool re-distributed efficiently to LDs when we blocked translation with cycloheximide 1h before LD induction by FA feeding (**Fig. S2H-J**).

### PLIN1 contains an unconventional integral membrane segment and a peripheral membrane binding motif

We next aimed at identifying the domains necessary for PLIN1 membrane-integration. We, therefore, carried out a systematic structure-function analysis using fluorescence microscopy and subcellular fractionation followed by alkaline extraction as parallel readouts.

We started with a minimal N-terminus of the mouse PLIN1 sequence (PLIN1 1-192) containing the PAT and 11-mer regions which are sufficient for LD targeting (**Fig. S3A**)^32,40,43^. As expected, neither of these two domains nor a polybasic region enriched in prolines downstream of the 11-mer region was involved in ER targeting. The N-terminal PLIN1 portion until position 237 remained completely soluble (**Fig. 3A**, S100) localizing to the cytoplasm and the endogenous LDs (**Fig. 3B, S3B**).

**Figure 3.**
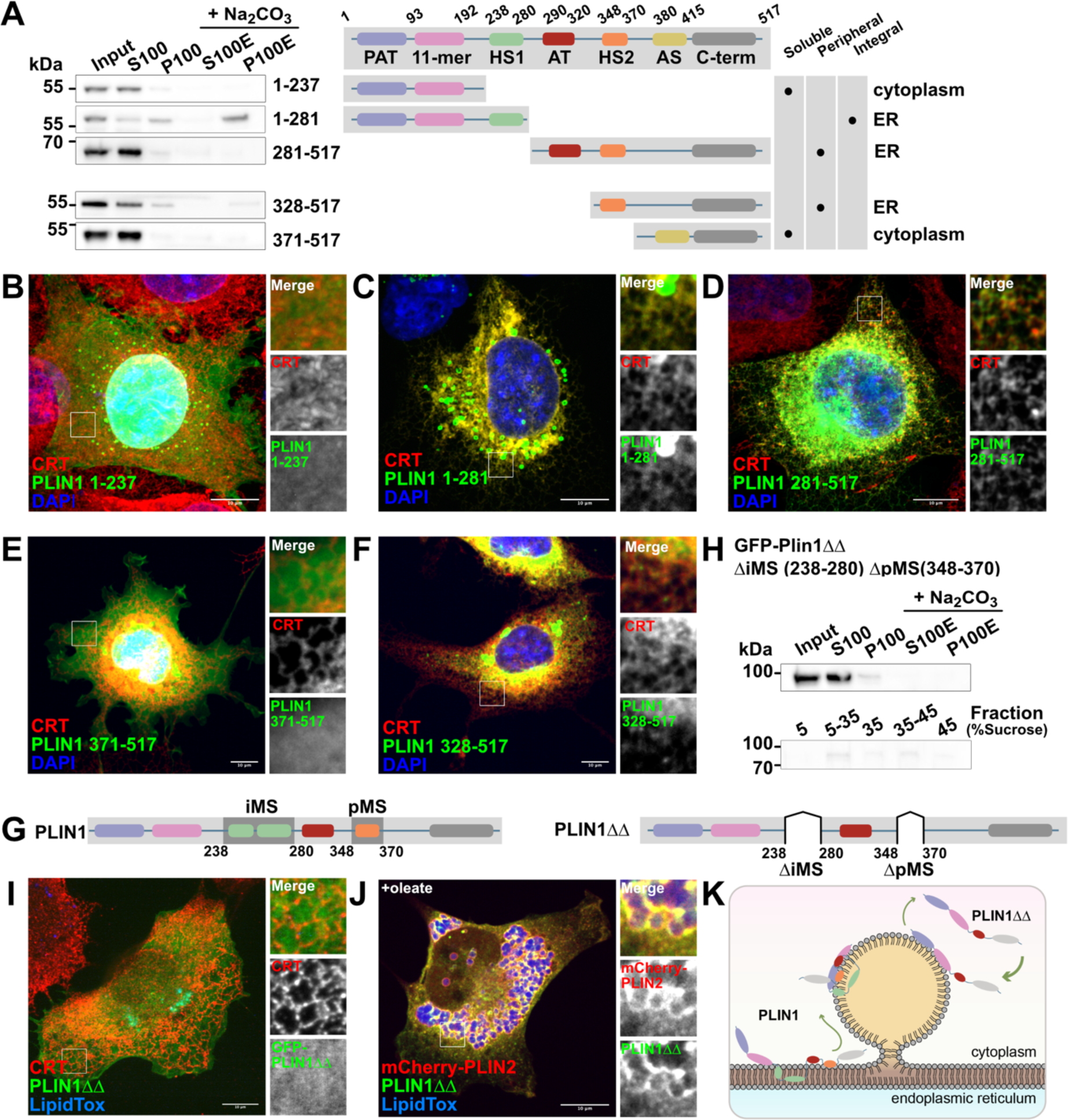
Identification of two ER-targeting segments in PLIN1. **(A)** PLIN1 analysis by fractionation and alkaline extraction and Western-blotting with anti GFP-antibody. See Fig.S3 for domain description. **(B)** COS7 cells expressing a GFP-PLIN1 (1-237). The ER was visualized with an antibody to calreticulin (CRT). The nuclei were stained with DAPI. Scale bar, 10 µm. **(C)** As in (B) but with PLIN1 1-281. **(D)** As in (B) but with PLIN1 281-519. **(E)** As in (D) but with PLIN1 371-517. **(F)** As in (D) but with PLIN1 328-517. **(G)** Illustration of the integral membrane segment (iMS, 238-280) and the peripheral membrane segment (pMS, 348-370) and PLIN1ΔΔ lacking both. **(H)** As in (A) but deletion of iMS and pMS (GFP-PLIN1ΔΔ). P100 was subjected to alkaline extraction (+Na_2_CO_3_) and floated as in Fig. 1G. **(I)** As in (B) but with PLIN1ΔΔ. LDs were stained with LipidToxDeepRed. **(J)** COS7 cells co-expressing m-cherry2-PLIN2, and GFP-PLIN1ΔΔ were treated with 250 µM Na^+^oleate for 6h. **(K)** Deletion of the iMS and pMS switches PLIN1 from an integral to class-II LD protein. Domain colors as in (A).

However, a construct extended until residue 281 (PLIN1 1-281) pelleted efficiently to the membrane fraction, was alkaline resistant (**Fig. 3A**, P100E), and localized to the ER as determined by microscopy, indicating that the region between residues 238 and 280 contained an integral membrane segment (iMS) (**Fig. 3A** green domain, and **S3B**). Interestingly, the corresponding C-terminal portion (PLIN1 281-517) was found in the supernatant S100 (**Fig. 3A**) but localized to the ER as shown by microscopy (**Fig. 3D**), suggesting that PLIN1 contains an additional peripheral membrane segment (pMS) contributing to ER-binding. Computational analysis of the C-terminal portion predicted a strong amphipathic region between residues 380 and 400 (**Fig. 3A**, yellow). However, a construct containing this motif (Plin1 371-517) remained completely soluble (**Fig. 3A, and 3E**). Further extension of the C-terminal part until the beginning of the 4-helix bundle (PLIN1 328-517) led to ER localization (**Fig. 3F**), but this construct remained soluble in the fractionation experiments (**Fig. 3A**), indicating that the potential pMS is positioned in this section of the middle domain (**Fig. 3A**, orange).

To narrow down the two membrane binding regions further, we turned to hydropathy plot analysis (**Fig. S2F**). A first segment with moderate hydrophobicity was found between tryptophan 238 and proline 280 of the mouse PLIN1 sequence, corresponding to the region in which we identified the iMS (**Fig. S2F and 3G**). A second slightly hydrophobic region between residues 348 and 370 overlapped with the pMS (**Fig. S2F and 3G**).

To test the function of these domains in ER targeting we deleted them. PLIN1ΔΔ lacking both the iMS and the pMS (**Fig. 3G**, PLIN1 Δ238-280, Δ348-370) was completely soluble, and did not float (**Fig. 3H**). Additionally, this mutant localized to the cytoplasm (**Fig. 3I**). Although having lost its ability to bind the ER, PLIN1ΔΔ was still found on LDs after FA feeding, and now often mixed with PLIN2 (**Fig. 3J**). In conclusion, removal of the iMS and the pMS converted PLIN1 into a class II LD-protein, targeting LDs directly from the cytoplasm (**Fig. 3K**).

In agreement with this model, addition of the iMS to PLIN2, right downstream of the 11-mer region (PLIN2-iMS (insertion at position 251)), was sufficient for membrane integration (**Fig S3C, S3D, S3E**). ER-insertion of PLIN2-iMS permitted efficient competition with PLIN1 for LD formation at the ER (**Fig. S3E**). However, in contrast to our expectation PLIN2-iMS did not mix with PLIN1 on the same LDs, but tended to segregate onto a separate LD class (**Fig. S3E**), indicating a critical function for the iMS in the nucleation of different LD populations at the ER.

Taken together our data show that PLIN1 behaves as an integral ER membrane protein showing characteristics of a class-I rather than a class-II LD-protein. Both the iMS and the pMS are sufficient for ER targeting, but membrane integration, and the generation of different LD-classes depend on the iMS. Removing the ability to integrate into the ER converts PLIN1 into a conventional class II LD-protein. The ER-binding deficient PLIN1-mutant mixes better with class II-PLINs on the same LDs indicating that it has reduced LD affinity.

### The iMS is critical in determining PLIN1 LD-binding kinetics

To investigate this possibility, we analysed the monolayer affinity of PLIN1 and PLIN1ΔΔ *in vitro* (**Fig. 4A**,^31^). Shrinkage of the aqueous compartments in a buffer-in-oil system leads to compression of an artificial LD-monolayer at the oil-buffer interface. The proteins fall off the monolayer with progressive reduction of the interface area and accumulate proportionally to their off-rates in the buffer compartment^31^. In agreement with the microscopy and fractionation experiments (**Fig. 3G-K**, and ^40,42^), full-length PLIN1 remained efficiently bound to the interface of the buffer compartment, indicating that the high LD affinity is determined by low off-rates from the monolayer (**Fig. 4, B**). In comparison, PLIN1ΔΔ was readily released from the interface, showing faster off-rates and thus lower affinity (**Fig. 4A, B**). The progressive accumulation in the buffer phase showed that PLIN1 excluded PLIN1ΔΔ from the LD surface mimic, confirming the results in cells (**Fig. 4A, B**).

**Figure 4.**
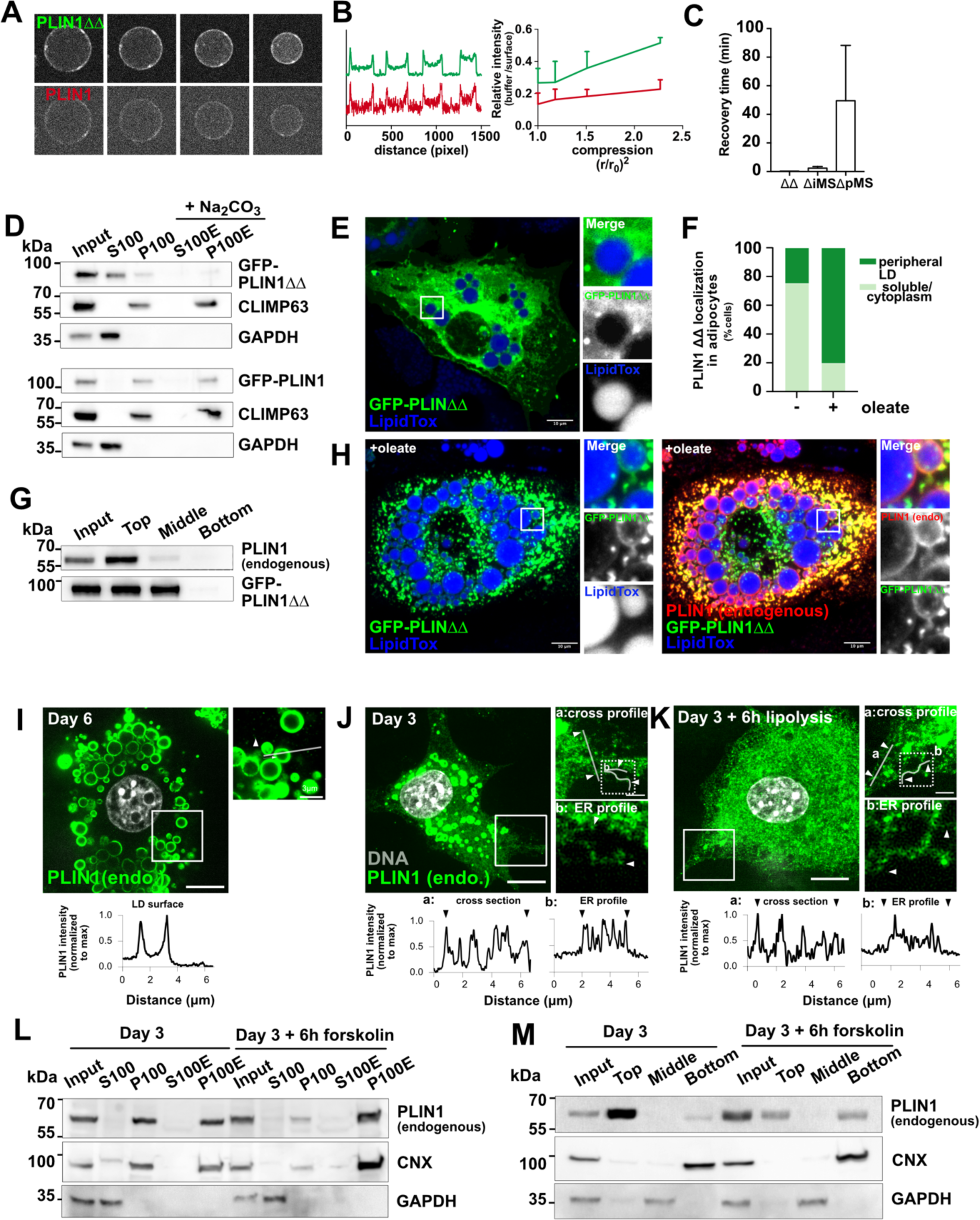
Membrane insertion determines LD-class specificity by tuning LD binding kinetics. (A) Localization of GFP-PLIN1ΔΔ and mCherry-PLIN1 in an buffer-in oil system with shrinking aqueous phase progressively shrinks. (B) Intensity ration of signal at the oil-buffer interface and within buffer for indicated compression ratios; GFP-PLIN1ΔΔ (green), mCherry-PLIN1 (red). N=3 independent experiments; mean ± SD. (C) FRAP with oleate treated Huh7 cells expressing GFP-PLIN1ΔΔ, ΔiMS or ΔpMS. Scale bar, 10 µm. N = 3 experiments; results are expressed as mean ± SD. (D) GFP-PLIN1ΔΔ or GFP-PLIN1 were expressed in 3T3-L1 cells from inducible transgenes and subjected to fractionation and alkaline extraction. Labeling as in Fig. 2; **CLIMP-63,** integral ER-membrane protein. (E) As in (D) but with adipocytes. PLIN1ΔΔ was induced 6 days post differentiation with 500 ng/ml anhydrotetracycline for 6h. LDs stained with LipidToxDeepRed. Scale bar, 10 µm. (F) Percentage of cells in (E) and (H) with GFP-PLIN1ΔΔ on peripheral LDs or in the cytoplasm, N = 100 cells. (G) As in (E) but with flotation. Endogenous PLIN1 and GFP-PLIN1ΔΔ were both detected with an anti PLIN1-antibody. (H) As in (E) but with oleate for 6h. GFP-PLIN1ΔΔ is now found on peripheral LDs. PLIN1ΔΔ remains excluded from the LDs covered by endogenous PLIN1. On the right, anti PLIN1-antibody stains both endogenous PLIN1 on large LDs (red) and PLIN1ΔΔ on the peripheral LDs (GFP+red=yellow). (I) Endogenous PLIN1 (green) in day6 adipocytes; line plot of PLIN1 intensity (normalized to maximum), as drawn in the inset. Scale bar: 10 μm, for inset 4 μm. (J) As in (I), but at day 3. Arrowheads mark terminal points of ER. (K) As in (J), but after addition of 10 µM forskolin for 6h (lipolysis). (L) Fractionation and alkaline extraction of adipocytes as in (J) and (K); labels as in (D); **CNX**, calnexin (integral ER-membrane protein). (M) Flotation experiment of the cells as in (J) and (K), done as in (G).

To gain insights into the corresponding on-reaction of the binding equilibrium, we used fluorescence recovery after photobleaching. Deletion of either the iMS (PLIN1ΔiMS) alone or together with the pMS (PLIN1ΔΔ) made recovery on the LD faster by more than an order of magnitude compared to a mutant in which the iMS is present and only the pMS is removed (PLIN1ΔpMS) (**Fig. 4C**, still frames in **Fig. S4A**).

Together these results suggest that PLIN1 binds LDs with low off and on rates, which is determined by the iMS. LD-binding via the classical amphipathic segments appears to be faster but shows lower LD-affinity.

These data suggested that differences in membrane targeting and LD-affinity by the PLINs may provide part of the mechanisms by LD-heterogeneity arises in adipocytes: PLIN1 is targeted to the ER via the iMS where it binds to the emerging LDs with high affinity excluding the low-affinity PLINs. However, because PLIN1 is kinetically trapped on this first LD population it cannot immediately bind to the second LD-class which emerges upon FA feeding. At this stage PLIN2, 3 and 4 are can readily bind to the surface of the acutely synthesized LDs from the cytoplasm. Despite their lower LD-affinity, the class-II PLINs label the second LD-population first.

### Perturbation of PLIN1 membrane targeting leads to a LD-class switch in adipocytes

To test these interpretations directly in adipocytes, we stably integrated GFP-PLIN1 and GFP-PLIN1ΔΔ as inducible transgenes into 3T3L1-preadipocytes. Consistent with the results above, PLIN1ΔΔ was soluble (**Fig. 4D**), and full-length PLIN1 behaved like an integral membrane protein (**Fig. 4D and 4E**). PLIN1 localized to LDs after FA feeding and as in COS7 cells moved from the ER to LDs even when translation was shut off with cycloheximide before LD induction (**Fig. S4B-E**).

After expression of the transgene in day 6 adipocytes, PLIN1ΔΔ behaved like a class-II LD protein being largely excluded from the big LDs which formed during differentiation (**Fig. 4E and F**). Endogenous full-length PLIN1 was expressed and floated with the LDs to the top fraction (**Fig. 4G**). PLIN1ΔΔ, was additionally found in the soluble middle fraction, confirming the microscopy experiments (**Fig. 4E and F**). Although being effectively excluded from the LDs covered by endogenous PLIN1 in cells, PLIN1ΔΔ remains LD binding competent (**Fig. 3J**, **Fig. 4A**), and likely gets access to TAGs when LDs break during mechanical cell-lysis (**Fig. 4G**).

Importantly, PLIN1ΔΔ efficiently moved from the cytoplasm to the peripheral LD population which forms after FA feeding (**Fig. 4H and F**). Consistent with the results shown for PLIN2-4 in **Fig. 1**, PLIN1ΔΔ remained largely excluded from the LDs that were covered by endogenously expressed full-length PLIN1 (**Fig. 4H and F**).

Taken together, these data suggest that removing the iMS and pMS not only reduced LD affinity and turned PLIN1ΔΔ into a class-II LD-protein, but also leads to a change in LD-class specificity (**Fig. S4F**).

Finally, we examined the localization of endogenously expressed PLIN1 during adipocyte differentiation. As expected, in day 6 adipocytes, PLIN1 localized almost exclusively to the surface of LDs (**Fig. 4I**). However, at earlier time-points, e.g. day 3, endogenous PLIN1 also abundantly present on the peripheral ER (**Fig. 4J and^10,44,45^**), and was completely alkaline-resistant confirming its membrane integration (**Fig. 4L)**. At this stage of differentiation, PLIN1 is probably stock-piled in the ER facilitating rapid expansion of the LD-surface.

Interestingly, when we triggered the breakdown of the LDs by activating lipolysis, PLIN1 efficiently moved at these stages back to the ER (**Fig. 4K-L**). The movement between LDs and the ER was further supported by flotation experiments, showing that stimulation of lipolysis re-distributed PLIN1 away from the remaining floating LDs into the membrane pellet (**Fig. 4M**).

## Discussion

Here, we have shown that contrary to its previous classification PLIN1 behaves as an integral membrane protein and is not classical class-II LD protein. We reveal that an unconventional integral membrane segment (iMS) likely folding as a hairpin is necessary and sufficient for integration into the ER membrane and facilitates high affinity binding to the LD.

PLIN1 moves onto the LDs as soon as they appear during adipocyte differentiation. Due to its high affinity, it excludes the other PLINs from these LDs already at earliest stages in the biosynthesis process. Up until differentiation day 3 the ER and LD pools readily exchange. However, at later stages (starting from day 6), PLIN1 is kinetically “trapped” on the surface of the first LD-class. As a result, the second LD-class which emerges upon FA feeding gets covered by PLIN2-4 which are class-II PLINs targeting the second LD-population from the cytoplasm.

Although we show with translation shut-off experiments that PLIN1 moves from the ER to the LD surface, it remains to be determined whether PLIN1 uses the conventional ERTOLD pathway (**Fig. S4F, ER→LD)**. The length of the iMS of around 30 amino acids (**Fig. S4G**) fits the size of known hairpin motifs, but the iMS exhibits surprisingly low hydrophobicity and does, therefore, not satisfy the basic definition of a conventional ER membrane anchor^46–48^. The iMS contains some features that are also found in other hairpins that move between the ER and the LDs^13^, such as prolines and large tryptophanes at the centre of the motif (**Fig. S4G**). Additionally, the iMS is flanked by positively charged residues (**Fig. S4G**)^13^, perhaps ensuring the correct membrane topology or contributing to the regulation of LD access and stability on the LD surface.

Consistent with other work^49^ our data indicate that the iMS favours the LD surface over the ER bilayer likely explaining why in adipocytes the ER pool is minimal at steady state. Since PLIN1 targets LDs early during biogenesis, it probably enters through the Seipin complex^28,50^, but may later on perhaps switch to the bridge pathway that was recently identified^51^ in fly, bearing molecular similarity with ER-LD contact sites previously identified in 3T3-L1 cells^52^. Our alkaline extraction data and detailed structure function seem incompatible with a cytoplasmic targeting pathway (**Fig. S4F ER→cytoplasm→LD**), but do not fully exclude this possibility.

The PLIN1 covered LDs are essential for adipose tissue function: Loss of PLIN1 in mice and humans can cause lipodystrophy^4,53^ because increased basal adipose lipolysis prevents the efficient LDs expansion^54,55^. The affected patients are highly insulin-resistant and suffer from hypertriglyceridemia, cardiovascular problems and fatty liver disease^55,56^. Partial loss of PLIN1 or PLIN1 haploinsufficiency can, surprisingly, have beneficial effects on metabolic health^57^. According to recent reports some PLIN1 mutants in fact improve metabolic profiles and reduce risk for cardiovascular disease^58,59^. A controlled perturbation of PLIN1 membrane targeting might therefore open avenues for mechanism-based therapy of fatty liver disease, type-2 diabetes, and obesity.

### Limitations of this study

A limitation of our study is that adipocyte differentiation in tissue culture may not fully recapitulate real tissue. Future work will address the physiological function of the two LD-classes that form in tissue culture.

Another open question is how PLIN1 is targeted to the ER. The simplest model involves membrane-integration by Sec61^60^. However, since the hydrophobicity of the iMS is relatively low, the pathway may depend the endoplasmic reticulum membrane complex (EMC) or requires a chaperone such as GET3/TRC40^61^. It is further still a possibility that in adipocytes PLIN1 is first inserted into the LD surface and then moves to the ER. However, if this were true, the acutely forming smaller LDs would then be covered by PLIN1 and not by different PLIN-family members.

Lastly, it is likely that additional machinery is required to form the two LD populations. We currently speculate that PLIN1 transport to the LDs is gated. The involved machinery may depend on the Seipin complex which probably contributes to the enrichment of PLIN1 on the surface of the first LD class and restricts the movement back to the ER.

## Acknowledgements

We thank Pedro Carvalho (University of Oxford) for critical reading of the manuscript and Lucas Pelkmans (University of Zurich) for generous support with research infrastructure. A.R.T. was supported by **ANR-MOBIL** and **ANR-21-CE11-LIPRODYN**. R.W.K acknowledges support from the Swiss National Science Foundation (**SNF 31003A_159793**), the Helmut Horten Foundation, and E.P. Abraham Cephalosporin fund. This publication arises from research funded by the John Fell Oxford University Press Research Fund and a grant from Diabetes UK (**22/0006453)**. R.W.K. acknowledges support from the imaging facility at the Dunn School of Pathology, University of Oxford.

## Author contribution

**Conceptualization,** R.W.K; **Methodology,** M.M., R.W.K., A.R.T; **Formal analysis,** M.M., O.S., D.A., K.B., M.O, A.R.T, R.W.K.; **Investigation,** M.M., O.S., D.A., K.B., M.O, A.R.T, R.W.K.; **Resources,** A.R.T. and R.W.K; **Writing – Original draft,** R.W.K.; **Writing – Review & Editing,** M.M., O.S., A.R.T., R.W.K., **Visualization;** M.M., O.S., A.R.T., R.W.K.; **Supervision,** A.R.T. and R.W.K.; **Funding Acquisition;** A.R.T and R.W.K.

## Experimental procedures

### STAR Methods

#### Lead contact and materials availability

Further information and requests for resources and reagents should be directed to and will be fulfilled by the Lead Contact, Robin Klemm.

#### Data and code availability

Original/source data for in the paper is available upon request from the lead contact and in Supplementary Data.

### Method Details

#### Cell lines

COS-7 cells were a kind gift from Tom A. Rapoport (Harvard Medical School, Boston, USA). COS7 is a male African green monkey kidney fibroblast cell line. HEK293T cells were obtained from Dharmacon (#TLP5918). HEK293T is a female human embryonic kidney cell line. 3T3-L1 pre-adipocytes were from ATCC (CL-173). 3T3-L1 is a continuous sub-strain of male murine 3T3 (Swiss albino) cells, developed through clonal isolation.

#### Cell culture

COS-7 cells were maintained in Dulbeccos’s Modified Eagle Medium with 4.5 g/l glucose and L-glutamine (DMEM, Thermo Fischer Scientific, #41965062), supplemented with 10% fetal bovine serum (FBS; Sigma-Aldrich, #F7524). HEK293T cells were maintained in DMEM containing 10% FBS, 25 mM HEPES (Sigma-Aldrich, H4034), 1 mM sodium pyruvate (Thermo-Fisher), 2 mM glutamine (Sigma-Aldrich) and 100 µM non-essential amino acids (Thermo-Fisher). All cells were kept as sub-confluent cultures and were sub-cultured regularly. All cell lines were periodically tested for mycoplasma and grown according to ATCC guidelines. Cells were incubated at 37°C and 5% CO_2_.

#### Pre-adipocyte maintenance

3T3-L1 pre-adipocytes were obtained from ATCC (CL-173) and cultured in DMEM with 4.5 g/l glucose and L-glutamine (Thermo Fischer Scientific, #41965062), containing 10% calf serum (CS; Sigma-Aldrich, #12133C). Cells were grown as sub-confluent cultures in 10 cm tissue culture dishes (TPP) and sub-cultured regularly.

#### Adipocyte differentiation procedure

For the differentiation of 3T3-L1 pre-adipocytes into white adipocytes, the cells were first grown to confluence and kept as a confluent culture for 48 hours. For differentiation in 96-well format, cells were seeded at a density of 10,000 cells per well of a 96-well plate (Greiner).

Typically, the cells reached confluence the next day. After 48 hours of confluence differentiation was induced by the addition of adipocyte differentiation medium consisting of DMEM supplemented with 10% FBS, 172 nM bovine insulin (Sigma-Aldrich, #I0516), 500 µM 3-Isobutyl-1-methylxanthine (IBMX, Sigma-Aldrich #I5879), and 1 µM dexamethasone (Gbiosciences, #API-04). Cells were kept in differentiation medium for 48 hours. The differentiation medium was then replaced by DMEM containing 10% FBS and 172 nM bovine insulin. Cells were kept in the insulin medium for another 48 hours. At day four after induction of differentiation the insulin medium was replaced by DMEM containing 10% FBS. The medium was then refreshed every two days. Typically, cells were used for experiments at day 6 or day 8 after start of differentiation.

### Method details

#### Transfection and subcellular fractionation and alkaline extraction

1.1×10^6^ COS-7 cells were seeded on a 10 cm dish the one day before transfection. The next morning cells were transfected with 10 µg of the plasmid of interest using GeneJuice transfection reagent (Merck, #70967-3) according to the manufacturer’s instructions. After 6 hours of incubation at 37°C the medium was replaced with fresh DMEM containing 10% FBS. The cells were then incubated at 37°C/5% CO_2_ overnight. The next morning the cells were washed once with ice-cold PBS and then harvested in 1 mL PBS with a cell scraper. The cells were briefly pelleted at 2000 g for 5 min at 4°C, and the supernatant was discarded. The cells were resuspended in 1 mL PBS containing protease inhibitor cocktail (cOmplete™, Mini, EDTA-free Protease Inhibitor Cocktail; Merck, #11836153001) (PBSI) and lysed by 20 passages through a bead-homogenizer (Isobiotec, 16 µm bead). A small aliquot for subsequent analysis by SDS-PAGE was taken (“Input”) and the remaining lysate was clarified by centrifuging for 10 minutes at 4°C and 2000 g. The supernatant was transferred to a 1.5 mL ultracentrifuge tube (Beckman-Coulter, #357448) and the membranes were pelleted at 100,000 g for one hour at 4°C in a TLA 100.3 rotor (Beckman-Coulter). Afterwards the supernatant was taken off and stored at −20°C (“S100”) and the pellet was resuspended in 50 µL PBSI. A small aliquot of 25 µL was stored at −20°C for analysis by SDS-PAGE (“P100”) and the remaining 25 µL were subjected to alkaline extraction by adding 75 µL of 250 mM Na_2_CO_3_ (pH 11) (Sigma Aldrich, #A0634292404) for 15 min on ice. The sample was transferred to a 250 µL ultracentrifuge tube (5 x 20 mm Beckman-Coulter, #342630) and a 50 µL of 10% sucrose cushion was placed below the sample at the bottom of the tube. The samples were then centrifuged for 1 h at 100,000 g and 4°C in a TLS 55 rotor (Beckman-Coulter) with the appropriate adapters (#358614). The top 200 µL were carefully taken off and stored at −20°C (“S100E”) and the pellet was resuspended in 50 µL PBS and stored at −20°C (“P100E”). The samples were subsequently analyzed by SDS-PAGE and western blotting to PVDF membranes (see respective sections).

#### Denaturing SDS polyacrylamide gel electrophoresis (PAGE) and immunoblotting

SDS-PAGE analysis was carried out either with 10% gels (TGX FastCast acrylamide Kit 10%; Bio-Rad) or precast 4-15% gradient gels (Mini-PROTEAN TGX precast gel; Bio-Rad). The protein concentration was determined using the Pierce 660 nm protein assay (Thermo-Fisher). Samples were denatured by the addition of SDS sample buffer (62.5 mM Tris-HCl, pH 6.8, 10% glycerol, 2% SDS, 0.002% bromophenol blue, 5% β-mercaptoethanol) and boiled for 10 minutes at 95°C. For western blotting the proteins were transferred to polyvinylidene difluoride (PVDF) membranes using the Trans-Blot Turbo RTA Mini PVDF Transfer Kit (Bio-Rad) and the Trans-Blot Turbo Transfer System (Bio-Rad). Membranes were blocked for one hour in 3% BSA/TBST (20 mM Tris-HCl, pH 7.4, 137 mM NaCl, 0.1% Tween 20) at room temperature. The incubation with the primary antibody was carried out in 3% BSA/TBST either for 1h at room temperature or overnight at 4°C. After extensive washing with TBST the membranes were incubated with the secondary HRP-conjugated antibodies diluted in 5% skim milk/TBST for 1h at room temperature. The membranes were again extensively washed with TBST and the ECL reagent (GE Healthcare) was mixed and added for 1 min afterwards. Images were acquired using Vilber-Lourmat instrument and its software.

#### Lipid droplet flotation

One 10 cm dish of day 6 3T3-L1 adipocytes was used. The cells were washed with ice-cold PBS and subsequently harvested by scraping in PBS. The cells were centrifuged at 600 g, 4°C for 5 minutes. The supernatant was discarded, and the pellet containing the cells was resuspended in 1 mL PBS supplemented with protease inhibitor (cOmplete™, Mini, EDTA-free Protease Inhibitor Cocktail; Merck). The cells were lysed by 20 passages through a bead homogenizer (16 µm bead, Isobiotec) and the lysate was clarified at 600 g, 4°C for 10 minutes. The supernatant was transferred to a new 1,5 mL tube and supplemented with 60% sucrose/PBS to a final concentration of 12% sucrose. 600 µL of the lysate were transferred to an ultracentrifuge tube (5 x 41 mm, Beckman-Coulter, #344090) and centrifuged in an MLS-50 rotor (Beckman-Coulter) at 100,000 g, 4°C for 1h. After the centrifugation the tubes were sealed, and the lower boundary of the floating lipid droplet fraction was marked. The tubes were flash frozen in liquid nitrogen and the top and were collected by cutting the frozen tube. The middle fraction was collected after thawing and the remaining pelleted fraction was washed once with PBS and then resuspended in PBS. All samples were then stored at −20°C until further analysis by SDS-PAGE.

#### Membrane flotation

Flotation experiments were carried out using five 10 cm dishes COS7 cells. Cells were washed with ice-cold PBS and subsequently harvested by scraping in PBS. The cells were briefly pelleted at 600 g, 4°C for 5 minutes. The supernatant was discarded, and the pellet was resuspended in 1 mL PBS supplemented with protease inhibitor (cOmplete™, Mini, EDTA-free Protease Inhibitor Cocktail; Merck). The cells were lysed by approx. 20 passages through a bead homogenizer (Isobiotec, 16 µm bead) and the lysate was cleared from debris at 600 g, 4°C for 10 minutes. 100 µL supernatant was collected and stored at −20°C; the remaining supernatant was transferred to a 1.5 mL ultracentrifuge tube (Beckman-Coulter) and centrifuged in a TLA 100.3 rotor (Beckman-Coulter) at 100,000 g, 4°C for one hour. The supernatant was transferred to a new 1.5 mL tube and stored at −20°C for later analysis. The pellet was resuspended in 36 µL PBS supplemented with protease inhibitor and 10 µL were stored for later analysis at −20°C. The remaining 26 µL were supplemented with 150 µL of 60% sucrose (w/v). To remove the ribosomes from the membranes 24 µL puromycin (10 mg/mL) were added to a final concentration of 2,5 M puromycin and 45% sucrose. The reaction was incubated at RT for 30 minutes. 300 µL of 35% sucrose (w/v) were added to a 600 µL ultracentrifuge tube (Beckman-Coulter) and the resuspended pellet fraction in 45% sucrose (w/v) was loaded to the bottom of the tube. Finally, 150 µL of 5% sucrose (w/v) were pipetted on top, the interphases were marked, and the sample centrifuged in a MLS 50 rotor (Beckman-Coulter) at 150,000 g, 4°C for 3h. The tubes were subsequently frozen in liquid nitrogen and the fractions separated by cutting the tube at the interface marks using a clean scalpel. The fractions that were collected are the top fraction with 5% sucrose, the interphase between 5% and 35% sucrose, the 35% sucrose fraction, the interphase between 35% and 45% sucrose and the bottom fraction with 45% sucrose. The protein concentration of the fractions was determined, and equal amounts analyzed by SDS-PAGE.

#### Indirect immunofluorescence

For indirect immunofluorescence experiments cells were trypsinized and transferred either to Ø 18 mm cover glasses (No. 1; VWR) in 12-well tissue culture plates (TPP) or to 96-well plates (Greiner) and cultured using DMEM supplemented with 10% of the respective serum (calf serum CS for 3T3-L1 or FBS for COS7 and differentiated adipocytes). Differentiated adipocytes were cultured for at least two days after trypsinization to ensure proper settling of the cells. Cells were fixed with 4% paraformaldehyde (PFA; Electron Microscopy Sciences) at 37°C for 30 minutes. After washing three times with PBS, cells were permeabilized for 15 minutes either with 0.1% Triton-X100 in PBS, or, when PLIN2 or PLIN3 or 4 were stained in adipocytes, 0.2% saponin and 1% CS in PBS. Cells were again washed and incubated with primary antibodies in blocking solution (PBS supplemented with 1% CS) for 1 h at RT. After washing, the cells were incubated with fluorescently labeled secondary antibodies in blocking solution for 1h at RT. For lipid droplet visualization HCS LipidTOX™ Green/Red/Deep Red dyes (Thermo-Fisher, #H34475/#H34476/3H34477, respectively) were used at 1:500 dilution in PBS. Nuclei were counterstained using DAPI (Thermo-Fisher) at a concentration of 0.4 µL/mL for 10 minutes at RT. Cell outline staining, if applied, was done using Atto 490LS NHS-Ester (ATTO-TEC) at a dilution of 1:800,000 in carbonate buffer for five minutes at RT.

Cover glasses were mounted onto Superfrost Excell glass slides (Thermo-Fisher) using Fluoromount G (SouthernBiotech). Image acquisition was performed as described under “Microscopy”.

#### Determination of protein concentration

Protein concentration was determined in 96-well plates using the Pierce™ 660nm Protein Assay (Thermo-Fisher) according to the manufacturer’s manual. A standard-curve was produced with a BSA solution ranging from 50-2000 µg/mL.

#### Microscopy

Confocal fluorescent microscopy was done on a Nikon Eclipse Ti-E microscope with enhanced CSU-W1 spinning disk (Microlens-enhanced dual Nipkow disk confocal scanner, wide view type), a Nikon CFI PlanApo 100x oil immersion objective (NA 1.49), and a sCMOS PCO Edge 5.5 camera (PCO, 2560×2160 pixels).

For automated confocal microscopy a Yokogawa CellVoyager 7000 spinning disk confocal microscope with enhanced CSU-X1 spinning disk (Microlens-enhanced dual Nipkow disk confocal scanner, wide view type), a 40x air objective (NA 1,15) or 60x water immersion (NA 1,2), and three Neo sCMOS cameras (Andor, 2560×2160 pixels) were used. 108 sites were acquired per well of a 96-well plate with seven confocal planes of 1 µm thickness per site, which were maximum intensity projected before saving. The signals for UV (405 nm) and far-red (640 nm) as well as green (488 nm) and Atto490LS (675 nm) signals were acquired in dual-camera mode. Red signal (561 nm) was acquired separately.

Images for parts of Fig. 4, S4 and S2H-I were captured on an Olympus SpinSR10 spinning disc confocal system fitted with an Olympus IX-83 frame, a Yokogawa CSU-W1 SoRa super-resolution spinning disc module, a Photometrics Prime BSI camera and a 60× objective (1.5 NA, UPLAPOHR60x), using Olympus cellSens Dimension software. When indicated the images were post-processed using an OSR filter (standard) and deconvolved (constrained iterative, maximum likelihood, 5 iterations) using Olympus cellSens Dimension software (version 3.1.1).

#### Automated image analysis

The acquired images were analyzed using CellProfiler (https://cellprofiler.org/). When necessary the cell profiler pipelines were run on the high-performance cloud computing system “ScienceCloud” at the University of Zurich. Analysis pipelines were composed of generic or partially customized modules as described in (Battich, Stoeger, & Pelkmans, 2015). Individual cells were segmented with nuclei as primary objects that were identified based on DAPI signal, and cell outlines were determined from Atto490LS signal using iterative segmentation based on a watershed algorithm and related to nuclei as secondary objects. After cell segmentation the background of all stainings of interest was subtracted and single-cell based features like area, shape, intensities and textures were extracted using the modules “MeasureObjectAreaShape”, “MeasureObjectIntensity”, and “MeasureTexture”. Subsequent analysis was done with Matlab (Mathworks).

#### Single cell classification by support vector machines (SVMs)

Single cells were classified using user supervised machine learning with a custom Matlab script CellClassifier (Rämö, Sacher, Snijder, Begemann, & Pelkmans, 2009). The classifier is based on support vector machine algorithm (SVM). In general, the classifiers were trained to identify and remove cells that were miss-segmented using nuclear area, nuclear shape, intensity, and textures as training features. Cells which overlapped with the border of an individual imaging site were automatically identified and removed. Furthermore, experiment specific SVMs were trained using intensity, channel correlation and texture as training features.

#### Cloning of truncations and deletion mutants

All truncation- and deletion-mutants were cloned by restriction ligation into the mEGFP-C1 using *Hind*III and *Kpn*I restriction sites. The mEGFP-C1 plasmid was a kind gift from Michael Davidson (Addgene plasmid #54759).

#### Cloning of GFP-PLIN2-251InsIMs

Cloning of the PLIN2 domain swap mutant carrying the core iMS (PLIN1 amino acids 238-268) of PLIN1 was done by insertion of the iMS sequence into mEGFP-C1-PLIN2 after amino acid 251 of the PLIN2 sequence. The insertion was done using the Q5 Site-Directed Mutagenesis Kit (NEB) and design of respective primers using the NEBaseChanger online tool (http://nebasechanger.neb.com/).

#### Generation of stable cell line inducibly expressing GFP-PLIN1

The stable 3T3-L1 cells with inducibe GFP-PLIN1 or GFP-PLIN1ΔΔ (Δ238-280-Δ348-370), respective transgenes were generated by lentiviral gene transfer using the pINDUCER20 construct and Gateway™ cloning technology. The pInducer20 was a kind gift from Stephen Elledge (Addgene #44012). The respective transgenes were first sub-cloned by TOPO cloning into a Gateway™ entry vector (pENTR™/D-TOPO™; Thermo-Fisher) and subsequently recombined into the pINDUCER20 vector using the Gateway™ LR Clonase™ II Enzyme mix (Thermo-Fisher). Production of lentiviruses and transduction of cells was performed as described below.

The cell lines were subsequently established by selecting positively transduced cells using G418 selection (1 mg/mL) for several passages.

Expression of the respective transgene was usually induced with 500 ng/mL anhydrotetracycline (ATC) for the indicated time-frames.

#### Preparation of sodium oleate

To prepare a solution of sodium oleate, 30 mL of double-distilled water were warmed up in the microwave to approx. 40°C and 400 µL of 5 M NaOH were added. While constantly stirring, 400 µL of oleic acid (Sigma-Aldrich) were slowly added and the solution was stirred for another five minutes (stock concentration 41 mM). If micelles where still visible, additional NaOH was added until all micelles disappeared. The solution was sterile filtered through a 0.22 µm sterile filter, aliquoted and stored at −20°C. Using 147 µL of the solution per 25 mL of culture medium resulted in a final concentration of 250 µM.

#### Cell treatments

Lipolysis was induced in differentiating adipocytes by addition of 1 mM forskolin in the medium (or vehicle, DMSO) to the final concentration of 10μM, for 6 hours. Oleate uptake was done by adding sodium oleate to the medium in the indicated concentration for 6 hours. Protein synthesis was inhibited by incubating cells in the medium containing cycloheximide (or DMSO control) to final concentration of 100μg/ml for the indicated time.

#### Cover glasses acid wash

Cover glasses (Ø 18 mm; VWR No. 1) were placed in a ceramic rack and submerged in 2 M HCl for 15 minutes. The coverslips were rinsed under tap water for 30 minutes and then washed three times with double-distilled water. Finally, the coverslips were rinsed in ethanol, sterilized under UV-light for 30 minutes and stored in a closed container in a tissue culture hood.

#### Production of lentivirus for gene transfer

Lentivirus for gene transfer and integration of pINDUCER constructs was produced in HEK 293T cells (Dharmacon). The cells were cultured in DMEM supplemented with 10% FBS, 25 mM HEPES (Sigma-Aldrich, H4034), 1 mM sodium pyruvate (Thermo-Fisher), 2 mM glutamine (Sigma-Aldrich) and 100 µM non-essential amino acids (Thermo-Fisher) on 10 cm dishes until reaching approx. 70% confluency. The cells were transfected using the calcium phosphate method with the three components necessary for the lentiviral packaging: 10 µg of the pINDUCER expression vector containing the construct of interest, 6.5 µg psPAX2 (Addgene # 12260), the viral genome packaging plasmid, and 3.5 µg pMD2.G (Addgene # 12259), encoding for the viral envelope components were gently mixed with H_2_O to a total volume of 597 µl. This was mixed with 682 µl 2x HBSS (Hank’s Balanced Salt Solution), and 85 µl 2M CaCl_2_ were added followed by 30 min incubation at RT. The entire transfection mix was added dropwise to the cells and incubated for approx. 12h at 37°C, 5% CO_2_. After that the medium was replaced by DMEM supplemented with 5% FBS. The media containing the lentiviral particles was collected at 24, 48, and 60 hours after changing the medium. The individual fractions were sterile filtered through a 0.45 µm filter, pooled and stored at 4°C. The virus particles were then concentrated to approx. 200 µL final volume using an Amicon Ultra-15 centrifugal filter unit (Merck) with a molecular weight cut-off of 100 kDa (Amicon “Ultra – 15 centrifugal filter”; 100 kDa MW-cut off). The filtrate was brought to 1 mL final volume with DMEM supplemented with 10% CS and immediately used for transduction of 3T3-L1 cells as described below.

#### Transduction of 3T3-L1 preadipocytes with lentivirus

3T3-L1 cells were grown in 6-well plates until 50-60% confluency before the medium was replaced by the concentrated virus mixture supplemented with 8 µg/mL polybrene (Sigma-Aldrich, #107689). After overnight incubation the medium was replaced by DMEM supplemented with 10% CS. Cells were further selected or subjected to single clone dilution as described below.

#### Single cell classification by support vector machines (SVMs)

Single-cell data obtained after automated image analysis was further analyzed using CellClassifier (Rämö, Sacher, Snijder, Begemann, & Pelkmans, 2009), a classifier software based on SVMs for supervised machine learning. In general, for all datasets SVMs were trained to identify and remove cells that were miss-segmented based on features like nuclear area, nuclear shape, intensity, and texture features. Also, datasets were automatically cleansed of cells which touch the border of an individual imaging site. Furthermore, experiment specific SVMs were trained e. g. for identifying transfected cells or cells which show a specific phenotype.

#### In vitro experiments

The purification protocol for LDs from cells expressing fluorescently tagged LD proteins was the same as in Ajjaji et al. 2019. Briefly, cells from 5 x 15 cm dishes were harvested, washed once in ice-cold PBS, and lysed using a 30G needle in 1ml 20 mM Tris-EDTA buffer containing complete protease and phosphatase inhibitor tablets at pH 7.5. To isolate LDs, 1ml of respective cell lysates were mixed with 1 ml 60% sucrose in 20 mM Tris-EDTA buffer supplemented with protease inhibitors, overlaid with 20%, 10% and 0% in buffered sucrose on top of one another in 5 ml Ultra-Clear centrifuge tubes (Beckman). Gradients were centrifuged for 16 h at 100,000 x *g* and 4 °C, using an SW60 rotor in a Beckman L8-70 centrifuge, and 300 µL were collected from the top as the LD fraction.

In vitro experiments were performed in HKM buffer: 50 mM Hepes, 120 mM K^+^OAc, and 1 mM MgCl_2_ (in Milli-Q water) at pH 7.4. To create buffer-in-oil drops, a buffer-diluted LD fraction was mixed with an excess of triolein by vortexing, as previously done in Ajjaji et al. 2019. The shrinkage of the buffer compartment was based on water evaporation of the aqueous drops, with the proteins at their surface, during imaging for 10 to 15 min on PDMS-coated glass slides.

#### Fluorescence recovery after photo-bleaching FRAP experiments

For FRAP experiments, we bleach the signal on a collection of drops and monitor the signal recovery. The background signal, e.g. from the cytosol, is removed from the recorded signal, which was at the end normalized by intrinsic bleaching of non-bleached areas. We used GraphPad Prism to fit the FRAP recovery curves with a non-linear regression and the exponential « one-phase association model ». The characteristic recovery time is extracted from the model.

#### Transfections

When indicated Huh7 cells (60-70% confluence) were exposed for 1hr to 500 µM oleic acid coupled to BSA (1% v:v) to induce LD formation and then cells were transfected with 3 µg of plasmid DNA/mL using Polyethylenimine HCl MAX (Polysciences, Inc) following the manufacturer’s instructions.

### Quantification and statistical analysis

Number of samples and replicates in an experiment, as well as dispersion (standard deviation, SD, standard error of the mean, SEM), are reported for each figure in the respective legends. Quantification of experimental data are described in relevant Method sections.

## KEY RESOURCES TABLE

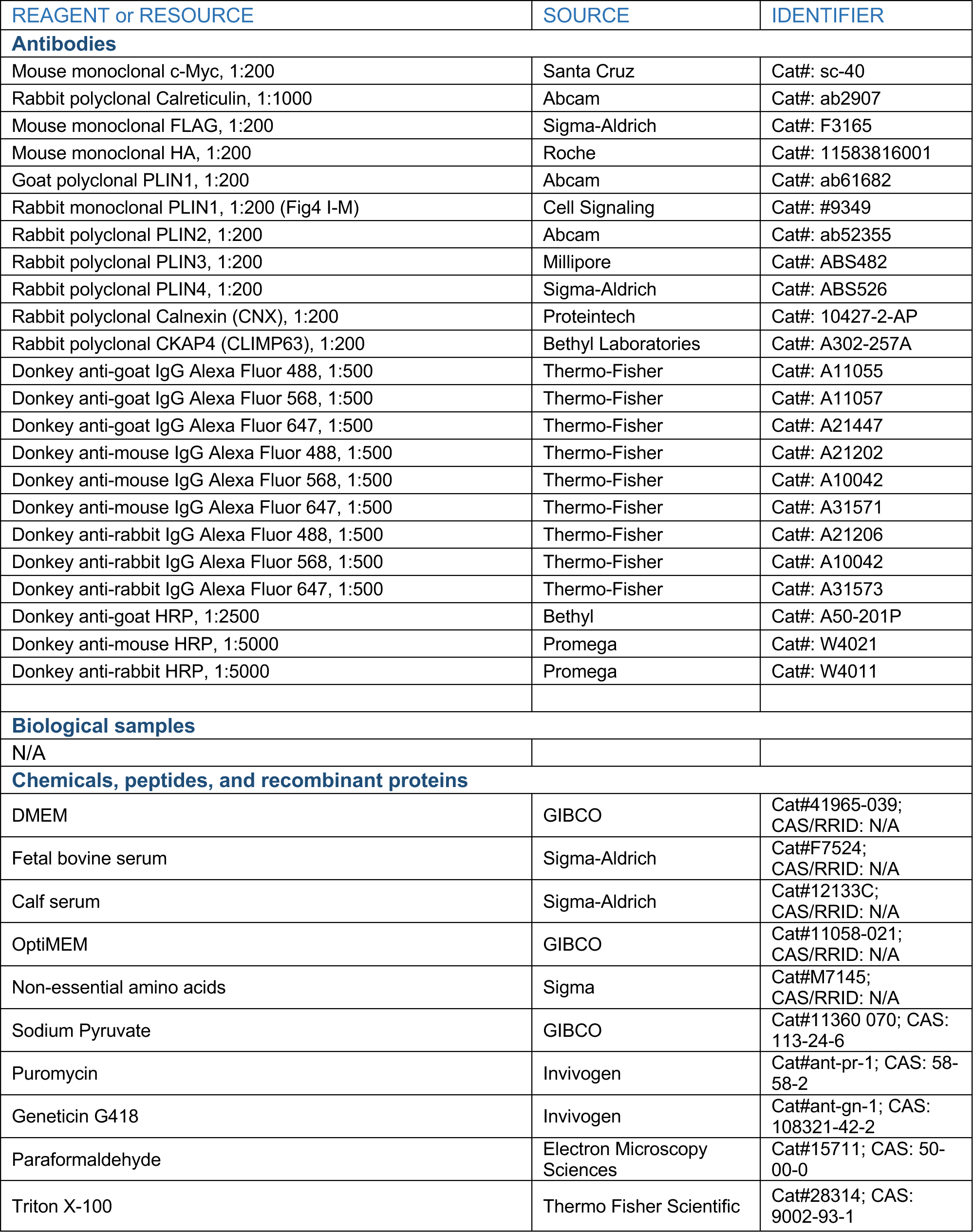
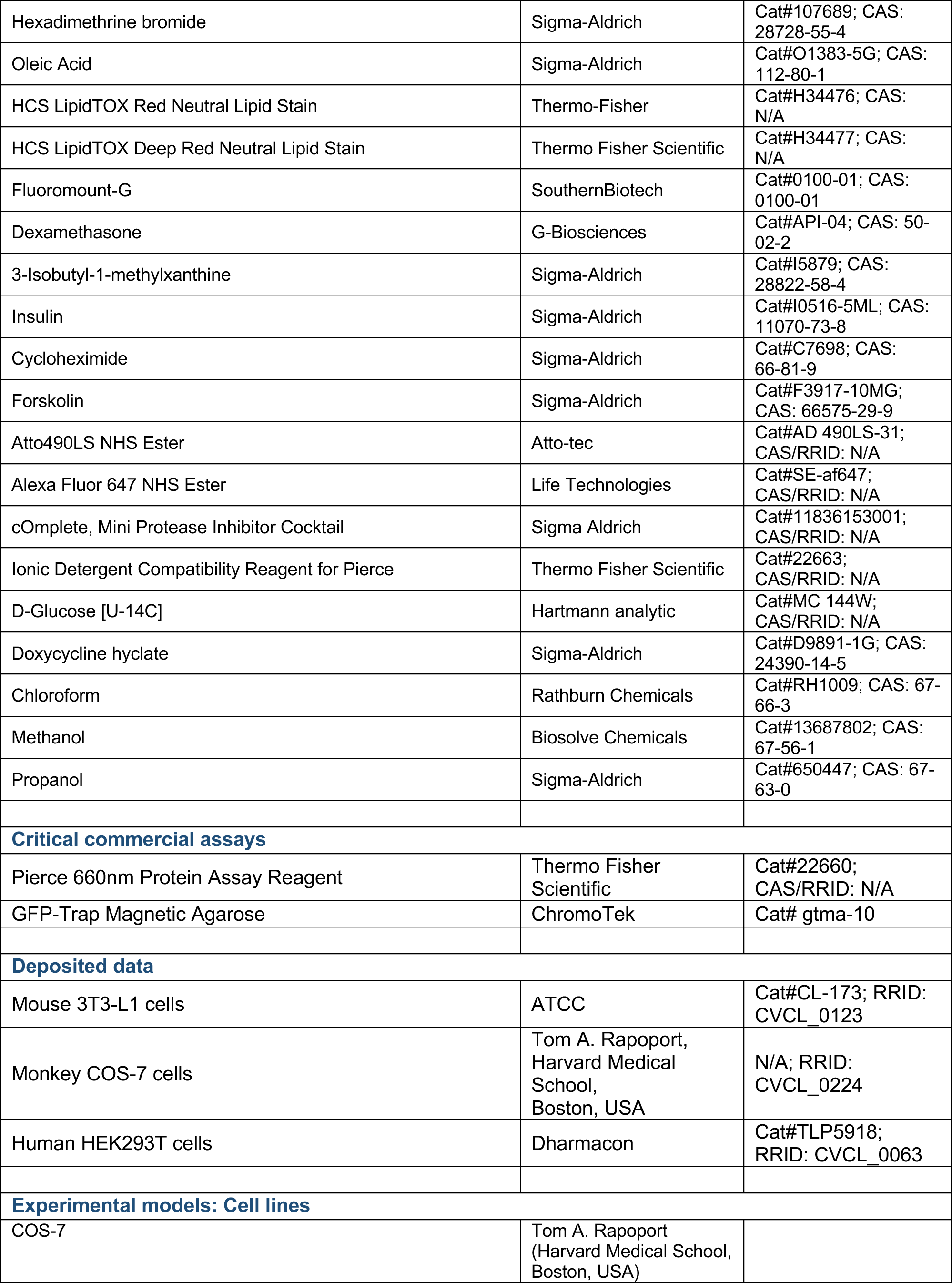
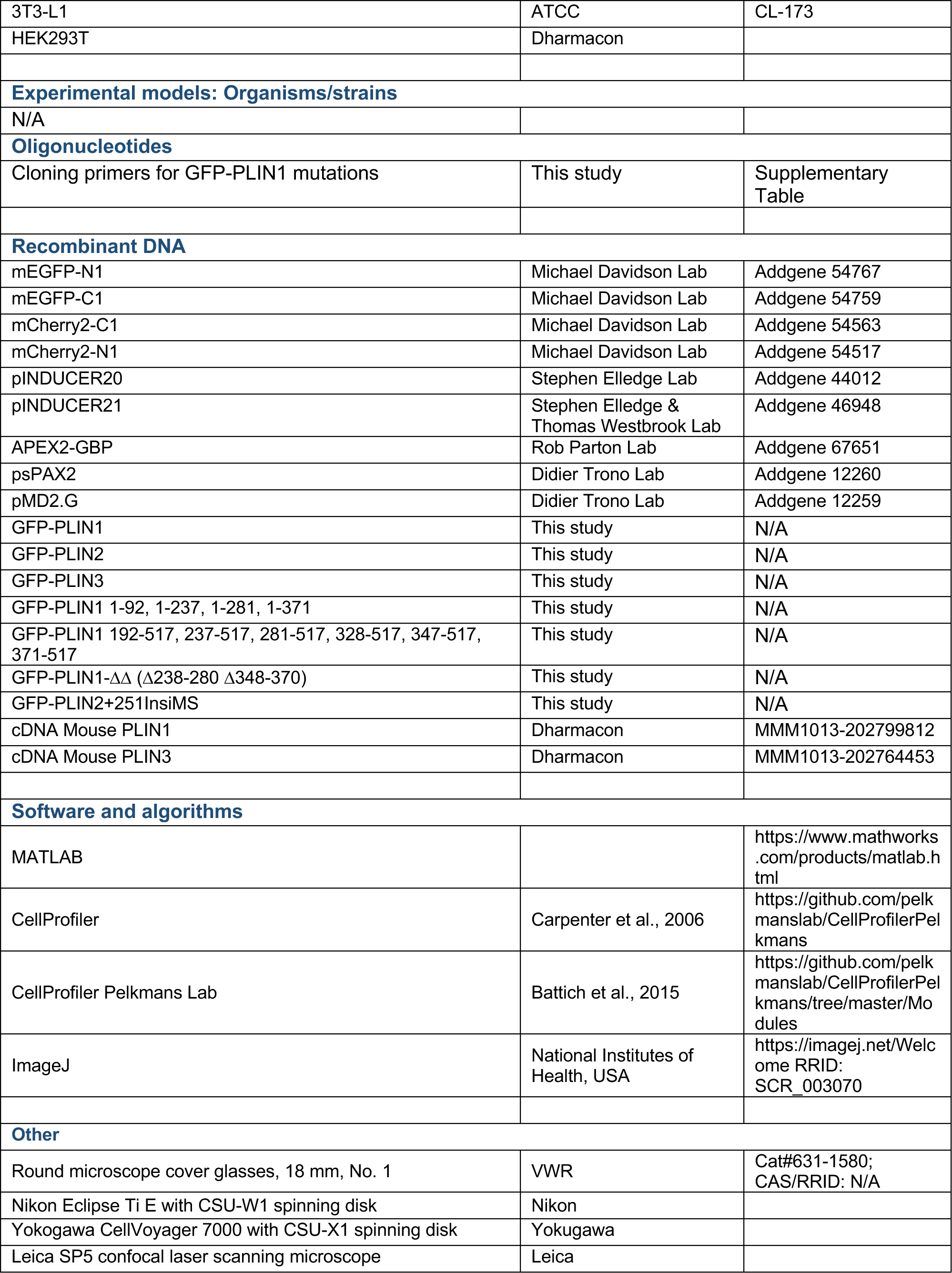

---

**Figure S1.**
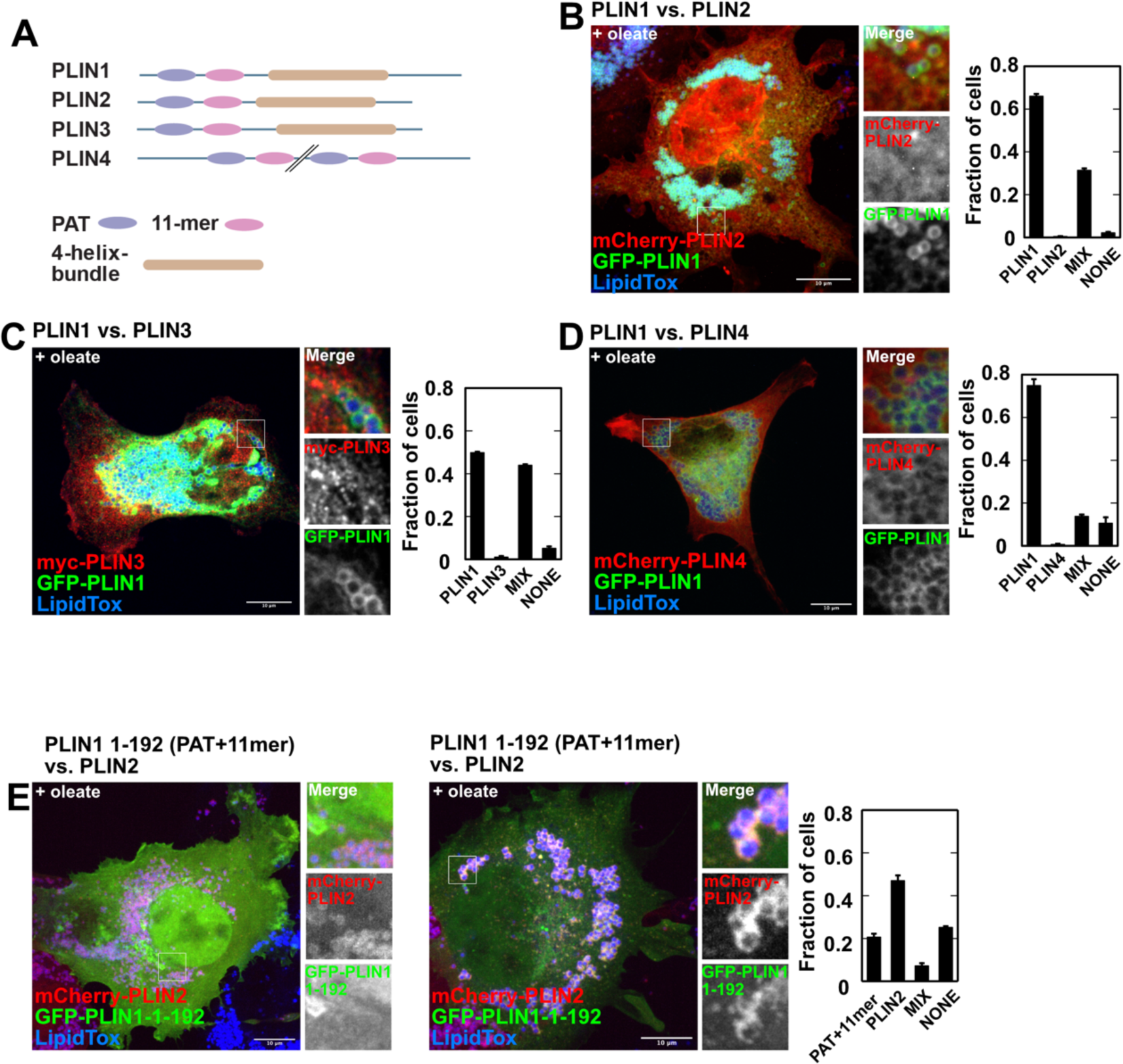
PLIN1 excludes PLIN2, 3 and 4 from LD binding in COS7 cells. **(A)** Schematic representation of perilipin (PLIN) 1-4. The N terminal PAT (perilipin, adipophilin/ADRP domain) and 11-mer regions are important for LD targeting. PLIN1-3 contain a predicted four helix bundle. The PLINs were grouped into a protein family based on the sequence similarity found in the PAT and 11-mer domains as well as their localization to the LD surface. PLIN4 does not contain a four-helix bundle and is much larger than the other family members. **(B)** COS7 cells co-expressing PLIN1 tagged at its N-terminus with GFP (GFP-PLIN1) and PLIN2 carrying an N-terminal mCherry2 tag (mCherry-PLIN2). 250 µM Na^+^oleate was added to the growth medium 6h prior to fixation. Lipid droplets were stained with LipidToxDeepRed (LipidTox). The cells were automatically segmented and assigned by a support vector machine based classifier into 4 groups, in which PLIN1 was exclusively found on the LD surface and PLIN2 was cytoplasmic **(PLIN1)**, cells that showed only PLIN2 LD staining **(PLIN2)**, cells in which various levels of PLIN2 was found on LDs together with PLIN1 **(MIX)**, and cells in which neither of the two proteins was found on the LD **(NONE)**. Intensity and texture of PLIN1 and PLIN2 channels as well as correlation with LipidTOXDeepRed were used as training features. n = 3 independent experiments; the results are expressed as mean ±SD. Scale bar, 10µm. **(C)** As in (B) but PLIN3 tagged at its N-terminus with c-myc (myc-PLIN3) was co-expressed with GFP-PLIN1. n = 3 independent experiments; the results are expressed as mean ±SD. **(D)** As in (C) but PLIN4 tagged at its N-terminus with mCherry2 (mCherry-PLIN4) was co-expressed with GFP-PLIN1. n = 3 independent experiments; the results are expressed as mean ±SD. **(E)** COS7 cells co-expressing PLIN1 truncated after the 11-mer at residue 192 tagged at its N-terminus with a GFP (GFP-PLIN1-1-192) and PLIN2 tagged at its N-terminus with mCherry2 (mCherry-PLIN2).

**Figure S2.**
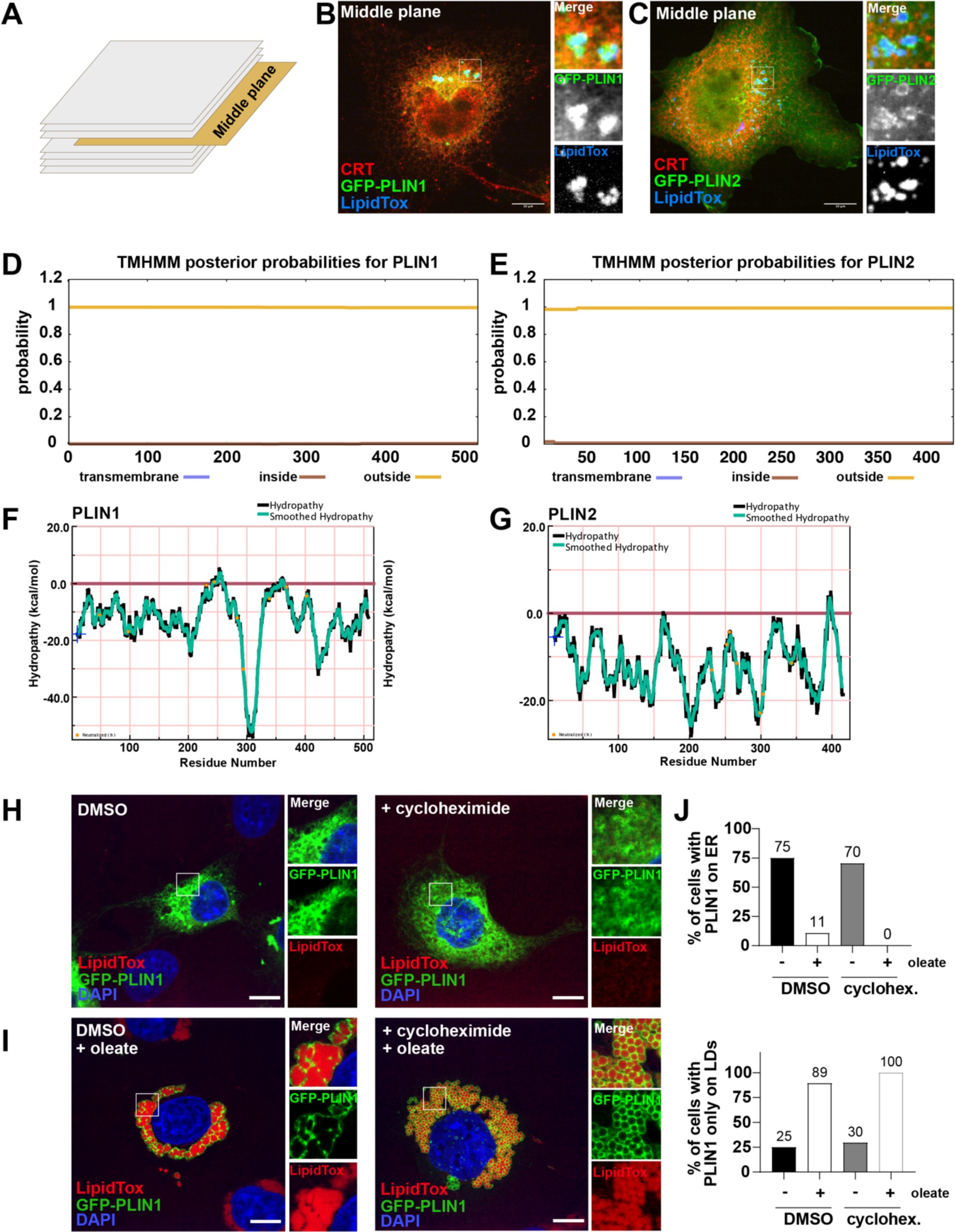
related to Figure 2 Subcellular targeting of PLIN1 and PLIN2; TMHMM predictions and hydropathy plot analysis for PLIN1 and PLIN2; PLIN1 LD targeting does not require translation. (A) Schematic illustration of image stack and the middle plane shown in (B) and (C). **(B)** COS7 cells expressing PLIN1 tagged at its N-terminus with GFP. Lipid droplets were stained with LipidToxRed and the ER was visualized with an antibody specific to the endogenously expressed luminal ER protein calreticulin (CRT). **(C)** As in (B) but cells expressed PLIN2 with an N-terminal GFP tag (GFP-PLIN2). **(D)** Topology analysis of PLIN1 using TMHMM 2.0 (Trans Membrane prediction Hidden Markov Model^62^ https://services.healthtech.dtu.dk/service.php?TMHMM-2.0). No integral membrane domains are predicted. **Outside,** segment in cytoplasm; **Inside,** segment inside the ER lumen; **transmembrane,** integral membrane segment. **(E)** As in (D) but for PLIN2. **(F)** Hydropathy plot analysis for PLIN1 with membrane protein explorer (MPEX)^63^. **(G)** As in (F) but for PLIN2. **(H)** COS7 cells expressing PLIN1 tagged at its N-terminus with GFP (GFP-PLIN1). Lipid droplets were stained with LipidToxRed (LipidTox) and nuclei were visualized with DAPI. Cells were either treated with DMSO as a control or with 100µg/ml cycloheximide 7h prior to fixation. Scale bar, 10 µm. **(I)** As in **(H)** but the cells were additionally treated with 200 µM Na^+^oleate for 6h prior to fixation (1h after addition of DMSO or 100µg/ml cycloheximide). The results are expressed as % of cells with GFP-PLIN1 on LDs. **(J)** Quantification of (H) and (I). The results are expressed as % of cells with GFP-PLIN1 on ER; DMSO: N=20 (-OA), N=28 (+OA); cycloheximide: N=27 (+OA), N=20 (-OA).

**Figure S3.**
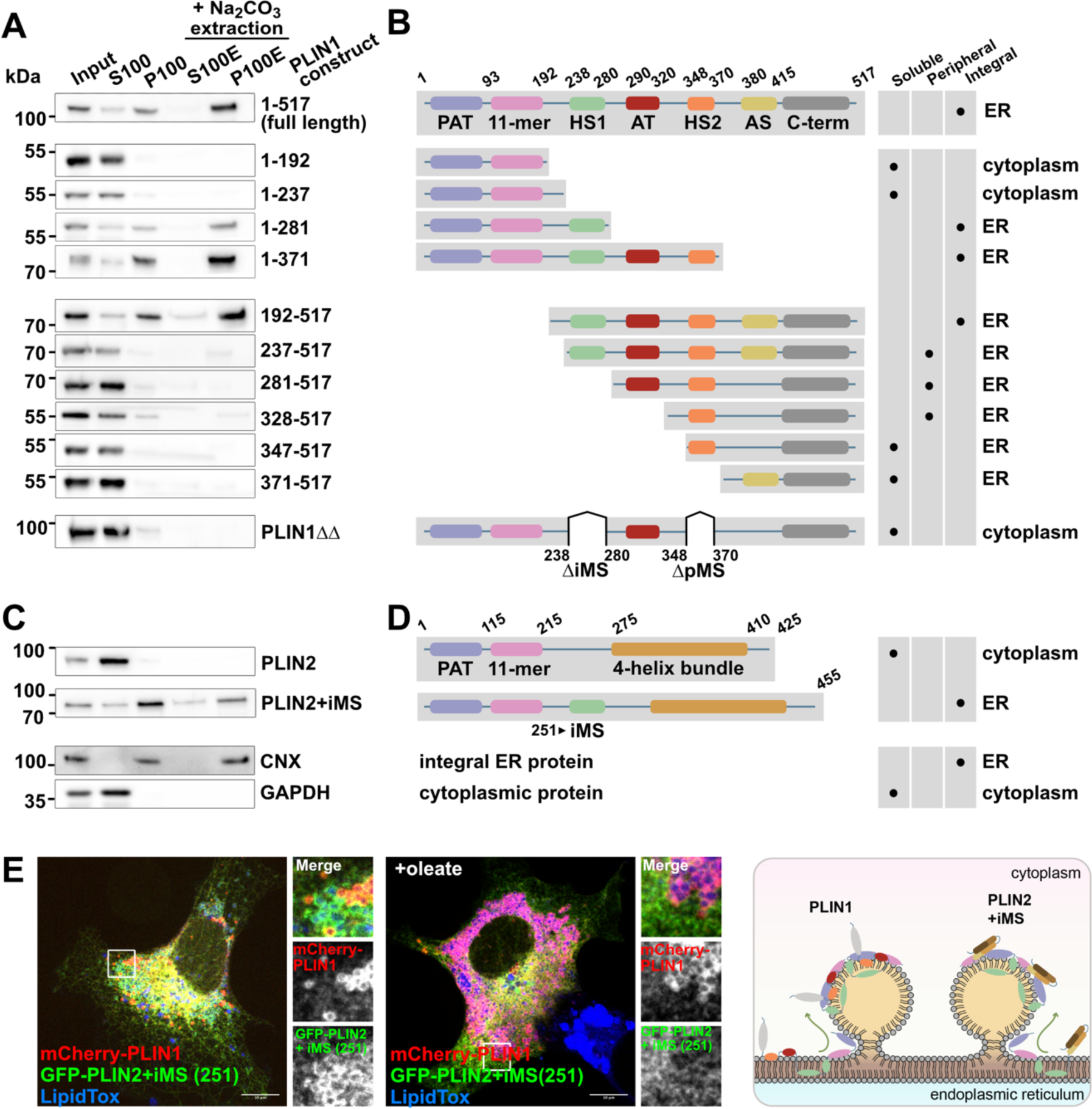
related to Figure 3 Summary of all constructs used in the structure function analysis of PLIN1. **(A)** Structure-function analysis of PLIN1 by fractionation and alkaline extraction using the indicated truncation constructs. **Input,** cell lysate; **S100, P100,** supernatant and pellet of a 100k x g centrifugation step, **S100E, P100E,** supernatant and pellet of 100k x g centrifugation after alkaline extraction with Na_2_CO_3_. Note that PLIN1 1-517 (full length) is also shown in Fig. 2F (GFP-PLIN1) and the constructs 1-237, 1-281 as well as 281-517, 328-517, and 371-517 are shown in main Fig. 3A. PLIN1ΔΔ is also shown in Fig. 3H. **(B)** Schematic illustration of the domains in PLIN1 and the tested truncation constructs. The table summarizes the results in (A) classifying the construct as soluble when found in S100 and in the cytoplasm by microscopy, as peripheral when found in S100, but seen on the ER by microscopy, and integral when present in the alkaline resistant membrane pellet P100E. ER, and cytoplasm is the localization as determined by microscopy. **PAT,** perilipin, adipohilin/ADRP, TIP-47 domain (purple); **11-mer,** LD targeting region (pink); **HS1, 2** hydrophobic segment 1 and 2 (green, orange, resp.); **AT,** acidic tract (red); **AS,** amphipathic segment (yellow); **C-term,** carboxy terminus with regulatory elements (grey). Classification of the constructs as soluble, cytoplasmic, peripheral or integral ER membrane proteins. **(C)** Fractionation and alkaline extraction experiments with GFP-PLIN2 (also shown in Fig 2F, for simplicity marked as PLIN2) and GFP-PLIN2 containing the iMS core from PLIN1 inserted after amino acid at position 251. For comparison with fractions in Fig. 2F the cytoplasmic marker **GAPDH,** glycerolaldehyde-3-P-dehydrogenase (from Fig. 2F) and the ER membrane protein marker **CNX,** calnexin (from Fig. 2F) are shown. **(D)** As in (B) but PLIN2 is shown. Classification and definitions as in (B). **(E)** COS7 cells were transfected with PLIN1 tagged at its N-terminus with mCherry2 (mCherry-PLIN1) and PLIN2 containing the iMS core from PLIN1 inserted after amino acid at position 251 (GFP-PLIN1+iMS (251). The cells were additionally treated with 250 µM Na^+^oleate 6h prior to fixation when indicated. The illustration shows that PLIN1 and PLIN2+iMs segregate preferentially to distinct LD classes showing that the iMS is critical in class specific LD nucleation at the ER. Domain colours are as indicated in (B) and (D).

**Figure S4.**
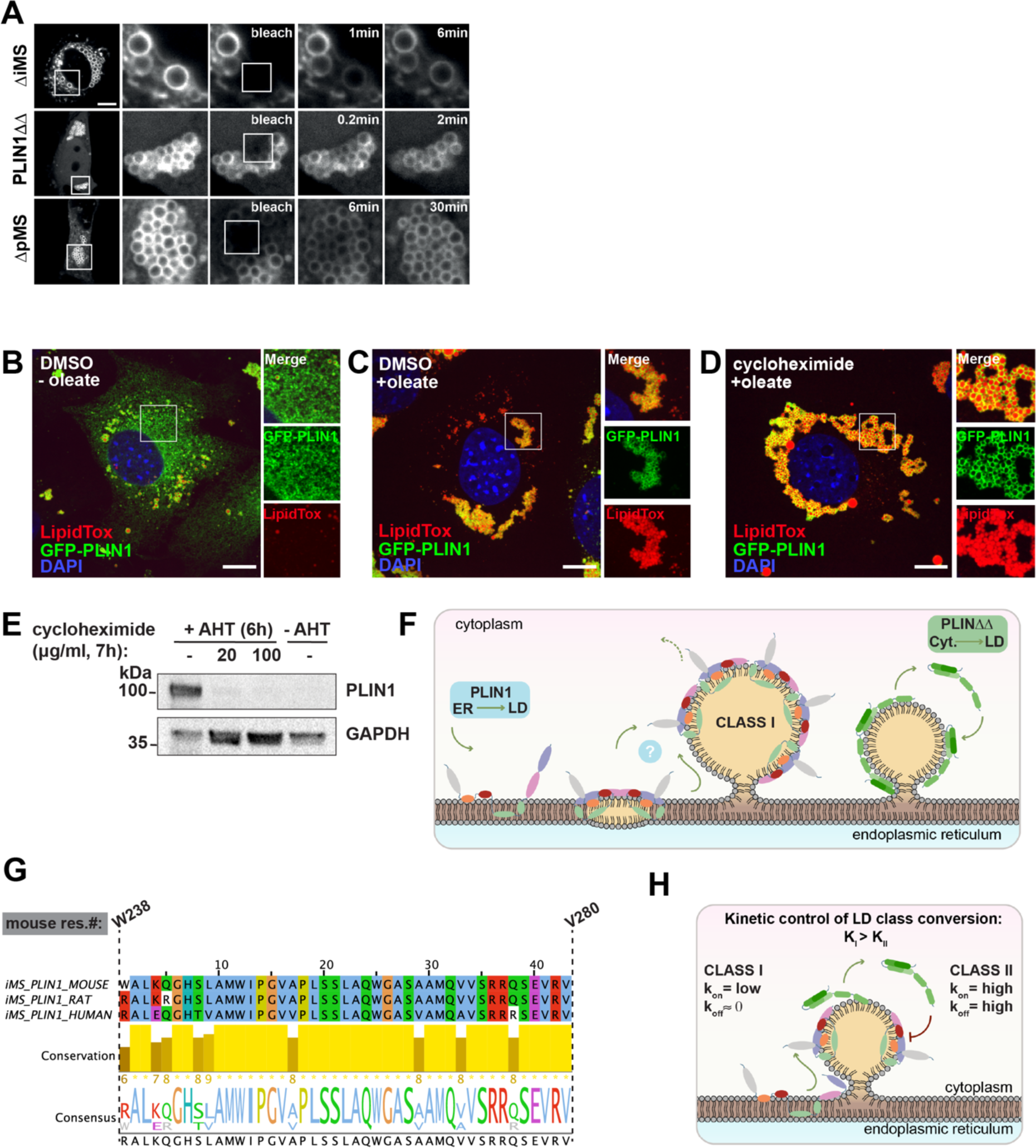
related to Figure 4 Recruitment of PLIN1 from the ER to the LD surface is independent of translation. **(A)** Huh7 cells expressing the indicated GFP-tagged PLIN1 constructs were subjected to photobleaching and the recovery of the GFP signal within the boxed area was plotted over time. **Related to Fig. 4C**. (B) 3T3-L1 cells expressing PLIN1 tagged at its N-terminus with GFP (GFP-PLIN1) from a stably integrated inducible trans-gene. Expression was induced with 500 ng/ml anhydrotetracycline (AHT) over-night (total 20h). Lipid droplets were stained with LipidToxRed (LipidTox) and nuclei were visualized with DAPI. Scale bar, 10 µm. **(C)** As in (B) but the cells were additionally treated with Na^+^oleate 3h prior to fixation. **(D)** As in (C) but translation was shut off by treating the cells with 100 µg/ml cycloheximide 1h before the addition of Na^+^oleate. **(E)** Validation of GFP-PLIN1 expression and translation shut off by cycloheximide. To induce expression of GFP-PLIN1, the cells were treated with 500ng/ml anhydrotetracycline (AHT) 6h prior to cell harvest. Translation was inhibited with the indicated concentrations of cycloheximide 1h before treatment with AHT. As a negative control AHT was omitted (-AHT). **(F)** PLIN1 is targeted to the ER membrane (brown) by the iMS (green) and the pMS (orange). PLIN1 likely moves to the LD using bridge-connections or by an unknown cytoplasmic intermediate that carries lipids to the LD (indicated by “?”). PLIN1ΔΔ binds to the LD directly from the cytoplasm as expected for a prototypical class-II protein. **(G)** Alignment of mouse, rat and human PLIN1 iMS. The regions correspond to mouse PLIN1 residues 238-280. The sequences were retrieved from Uniprot Q8CGN5, P43884, O60240 respectively, aligned using ClustalOmega https://www.ebi.ac.uk/Tools/msa/clustalo/ and analysed in JalView https://www.jalview.org/ using the T-coffee algorithm for determination of the consensus sequence shown below the sequence conservation bar graph. **(H)** LD-binding rates of class-II proteins are faster than for class-I proteins. The steady state PLIN1 ER-pool is low, slowing down the LD on-reaction. When bound to the LD, class-II proteins exchange faster than class-I proteins. The resulting high affinity to LDs leads to the gradual replacement of class-II by class-I proteins from the newly formed LDs.

